# Nanoscale dynamics of streptococcal adhesion to AGE-modified collagen

**DOI:** 10.1101/2022.10.23.513419

**Authors:** Camila Leiva-Sabadini, Paola Tiozzo-Lyon, Luis Hidalgo-Galleguillos, Lina Rivas, Agustín I Robles, Angélica Fierro, Nelson P Barrera, Laurent Bozec, Christina MAP Schuh, Sebastian Aguayo

## Abstract

The adhesion of initial colonizers such as *Streptococcus mutans* to collagen is critical for dentinal and root caries progression. One of the most described pathological and aging-associated changes in collagen – including dentinal collagen – is the generation of advanced glycation end-products (AGEs) such as methylglyoxal (MGO)-derived AGEs. Despite previous reports suggesting that AGEs alter bacterial adhesion to collagen, the biophysics driving oral streptococcal attachment to MGO-modified collagen remains largely understudied. Thus, the aim of this work was to unravel the dynamics of the initial adhesion of *S. mutans* to type-I collagen in the presence and absence of MGO-derived AGEs, by employing bacterial cell force-spectroscopy with atomic force microscopy (AFM). Type-I collagen gels were treated with 10mM MGO to induce AGE formation, which was characterized with microscopy and ELISA. Subsequently, AFM cantilevers were functionalized with living *S. mutans* UA 159 or *S. sanguinis* SK 36 cells and probed against collagen surfaces to obtain force-curves displaying bacterial attachment in real-time, from which the adhesion force, number of events, Poisson analysis, and contour and rupture lengths for each individual detachment event were computed. Furthermore, in-silico docking studies between the relevant *S. mutans* UA 159 collagen-binding protein SpaP and collagen were computed, in the presence and absence of MGO. Overall, results showed that MGO modification increased both the number and adhesion force of single-unbinding events between *S. mutans* and collagen, without altering the contour or rupture lengths. Both experimental and in-silico simulations suggest that this effect is due to increased specific and non-specific forces and interactions between *S. mutans* UA 159 and MGO-modified collagen substrates. In summary, these results suggest that collagen alterations due to glycation and AGE formation may play a role in early bacterial adherence to oral tissues, associated with conditions such as aging or chronic hyperglycemia, amongst others.

## Introduction

In recent years, populations worldwide have experienced a rapid increase in life expectancy that has brought forward many challenges in diagnostics and treatment for older people (Kontis et al. 2017). In dentistry, aging is associated with highly prevalent oral pathologies such as dental caries and especially root caries (Murray Thomson 2014). Once the root surface becomes exposed, demineralization of the superficial dentin layer due to diet-related pH changes, bacterial activity, or restorative dentistry procedures can occur and expose the underlying collagen matrix (Takahashi and Nyvad 2016). The collagen matrix becomes an essential substrate for streptococcal adhesion via specific surface adhesion proteins (i.e., adhesins) known as collagen-binding proteins (Cbps). Most notoriously, amongst initial colonizers *Streptococcus mutans* expresses clinically-relevant Cbps such as SpaP (Sullan et al. 2015; Avilés-Reyes et al. 2017), which are critical for dentinal caries progression in both animal and human studies as well as for invasion into tissues and the bloodstream for remote infections (Nomura et al. 2009). The adhesion of bacteria to surfaces is considered the initial factor for biofilm formation to take place and the physicochemical, mechanobiological, and molecular properties of the substrate play a key role in the initial bacterial attachment (Hojo et al. 2009; Berne et al. 2018). In dental biofilms, these initial surface colonizers are mostly comprised of streptococci, including *S. mutans* and *S. sanguinis*, amongst others (Yukari et al. 2021).

Nowadays new attention has been placed on investigating age-related changes in collagen, as it is one of the main structural components of dentin and periodontal tissues (Bailey et al. 1998; Gurav 2013). One of the most described aging-associated changes in collagen is non-enzymatic glycation, a process that leads to the generation of advanced glycation end-products (AGEs) (Nass et al. 2007), such as pentosidine and methyl-glyoxal (MGO) (Gkogkolou and Böhm 2012). MGO is derived from the metabolism of glucose and has the ability of attaching to lysine and arginine residues on collagen molecules (Wetzels et al. 2017), driving the formation of MGO-mediated adducts and crosslinks in a time-dependent manner. Thus, the accumulation of AGEs is higher in aged tissues, being most prominent in those with slow or negligible collagen turnaround, such as dentin (Semba et al. 2010; Ahmed et al. 2017). Although the effect of AGEs in modulating human cell attachment is well explored (Egaña-Gorroño et al. 2020), their role in altering bacterial adhesion and biofilm formation remains poorly understood. Recently published work suggested that MGO-derived AGE accumulation in an in-vitro collagen model increases the stiffness of collagen fibrils and enhances early adhesion of *S. mutans* over *S. sanguinis* (Schuh et al. 2020), implying that MGO-derived AGEs may potentially promote dysbiosis during early stages of collagen attachment with atomic force microscopy (AFM). However, the particular biophysics of bacterial-surface interactions remains unknown. Thus, the aim of this work was to unravel the dynamics of the initial adhesion of *S. mutans* to collagen in the presence and absence of MGO-derived AGEs, by employing bacterial cell force-spectroscopy with atomic force microscopy (AFM).

## Methods

### Type-I collagen gel fabrication

1mg/ml type-I collagen gels were made from rat tail collagen (Gibco) as previously described in a 96-well plate (Schuh et al. 2020). After a 24hr incubation, gels were incubated with 10mM MGO (Sigma) for 7 days at 37°C to induce MGO-AGE formation in the collagen matrix. Incubation with phosphate buffer saline (1x PBS) was used as a control.

### MGO-AGE quantification in the collagen matrix

Following incubation with MGO, AGE formation was assessed with ELISA and collagen autofluorescence. For ELISA, supernatants were removed from the wells and gels were snap-frozen on day 7 and stored at −80°C until further analysis. Gels were homogenized with micro-pistils and incubated with RIPA buffer (1x, Sigma) overnight. Gel remnants were removed by centrifugation (10 min, 1.000xg) and protein content was measured with BCA assay (ThermoFisher, US). Samples were normalized to 100μg/ml with ddH_2_O. The MGO-derived hydroimidazolone MG-H1 was measured using competitive ELISA (Abcam, UK) according to the manufacturer’s instructions. Briefly, protein binding 96-well plates were coated with conjugate in PBS and incubated overnight at 4°C. After blocking with assay diluent for 1 hour, wells were incubated with sample or assay standard (50 μl) for 10 min. Subsequently 1x anti-MG-H1 antibody (50μl) was added for 1 hour. Wells were washed and incubated with a secondary antibody (HRP-conjugated). The assay was developed with substrate solution (R&D systems) and the reaction was stopped with 1M H_2_SO_4_ and analyzed with a Tecan Sunrise microplate reader at (Tecan, Austria) at 450nm. For collagen autofluorescence, gels were prepared and incubated as above in black-sided 96-well plates for 7 days. Autofluorescence was measured at 360nm excitation and 460nm emission on days 0, 1, 4, and 7 using a multimodal microplate reader (Synergy HT, Biotek, USA).

### Scanning electron microscopy (SEM) and AFM topographical characterization of collagen gels

For SEM preparation, PBS- and MGO-treated collagen (MG-H1col) were fixed with 4% paraformaldehyde for 12 hours, followed by an ethanol series dehydration (25%, 50%, 75%, 90%, and three washes in 100% ethanol) and critical point drying in hexamethyldisiloxane. Samples were then sputter-coated with gold and observed with a JEOL JSM-IT300 LV SEM. Furthermore, AFM topographical images were obtained with individually tuned TAP300GD-G cantilevers (BudgetSensors, Bulgaria) in intermittent contact mode in ambient conditions, after gel physisorption onto glass cover slips. 2×2μm scans were performed on top of fibrillar collagen regions, and 3D reconstruction images were created using proprietary Asylum Research AFM software (v. 16.10.211).

### Bacterial probe functionalization and force-spectroscopy experiments

BL-TR400PB iDrive cantilevers (k~0.09N/m, Asylum Research, US) were functionalized by immersion in a 50μl droplet of 0.01M poly-L-lysine solution (PLL, Sigma, US) for 2 min. Excess PLL was rinsed off with ultrapure water and cantilevers were airdried for 10min. Subsequently, for the custom functionalization of bacterial AFM probes, *S. mutans* UA 159 and *S. sanguinis* SK 36 were employed. After 24hr growth at 37°C and 5% CO_2_, bacteria were adjusted to a concentration of ~1×10^8^ CFU/ml in 1x PBS, and gently vortexed for homogenization. A 50μl droplet of the bacterial solution was placed on the cantilever and incubated for 2 mins. Cantilevers were then gently washed with 1x PBS to remove non-adhered bacteria and immediately transferred to the AFM for experiments.

Single-cell force-spectroscopy measurements between functionalized bacterial probes and collagen substrates were carried out in a hydrated environment (PBS). In iDrive mode, bacterial probes were approached to the collagen surface with a maximum loading force of 0.2nN and a constant speed of 0.5Hz. A dwelling time of 5s was used to ensure appropriate adhesion between bacteria and collagen surfaces. Measurements were performed on five independent gels. The overall adhesion work and single-unbinding event rupture forces, rupture lengths, and contour lengths were computed from the recorded force-curves utilizing proprietary Asylum Research AFM software. Furthermore, a Poisson analysis was performed on the dataset to calculate short- and long-range force components from bacterial AFM unbinding experiments (Chen et al. 2011).

### Computational three-dimensional structures

The crystal structure of type-I collagen was obtained from the Protein Data Bank (PDBid: 7CWK). Once the structure was submitted to H++ server to add hydrogen atoms and to compute the pKa values of ionizable groups. Then, collagen modified with MGO to form MG-H1 was generated using the PyMOL program, and the Molecular sculpting function was used as a real-time energy minimizer (The PyMOL Molecular Graphics System, Version 2.0 Schrödinger, LLC.). The whole structure of SpaP protein was obtained using AlphaFold Protein Structure Database (https://alphafold.ebi.ac.uk/) which is based on artificial intelligence (AI) to predict the three-dimensional structures. Chimera (Pettersen et al. 2004) was used to add hydrogen atoms. To generate each protein-protein complex, docking studies were carried out using HDock server (Yan et al. 2020). A hybrid algorithm of template-based and template-free docking was used to generate the most probable protein-protein complexes as well as associated docking scores. The electrostatic potential studies were done using Adaptive Poisson-Boltzmann Solver (APBS) (Baker et al. 2001) in AutoDockTools.

### Statistical analysis

All resulting data were tabulated and analyzed using GraphPad Prism 9. After outlier detection, the dataset was tested for normal distribution. Statistical significances were assessed with t-tests or Kruskal-Wallis tests, considering a significance value of p<0.05.

## Results

### Collagen substrate characterization

1mg/ml collagen gels were fabricated according to previously published methods and incubated with 10mM MGO for 7 days to promote AGE formation. ELISA analysis of resulting gels confirmed the incorporation of the relevant MGO-derived hydroimidazolone MG-H1 AGE into collagen after 7 days, which was accompanied by a significant increase in collagen autofluorescence **(Figures 1A and 1B)**. SEM imaging displayed changes in fibril ultrastructure and morphology after 7-day MGO incubation, which was confirmed by AFM imaging **(Figure 1C)**.

**Figure 1:**
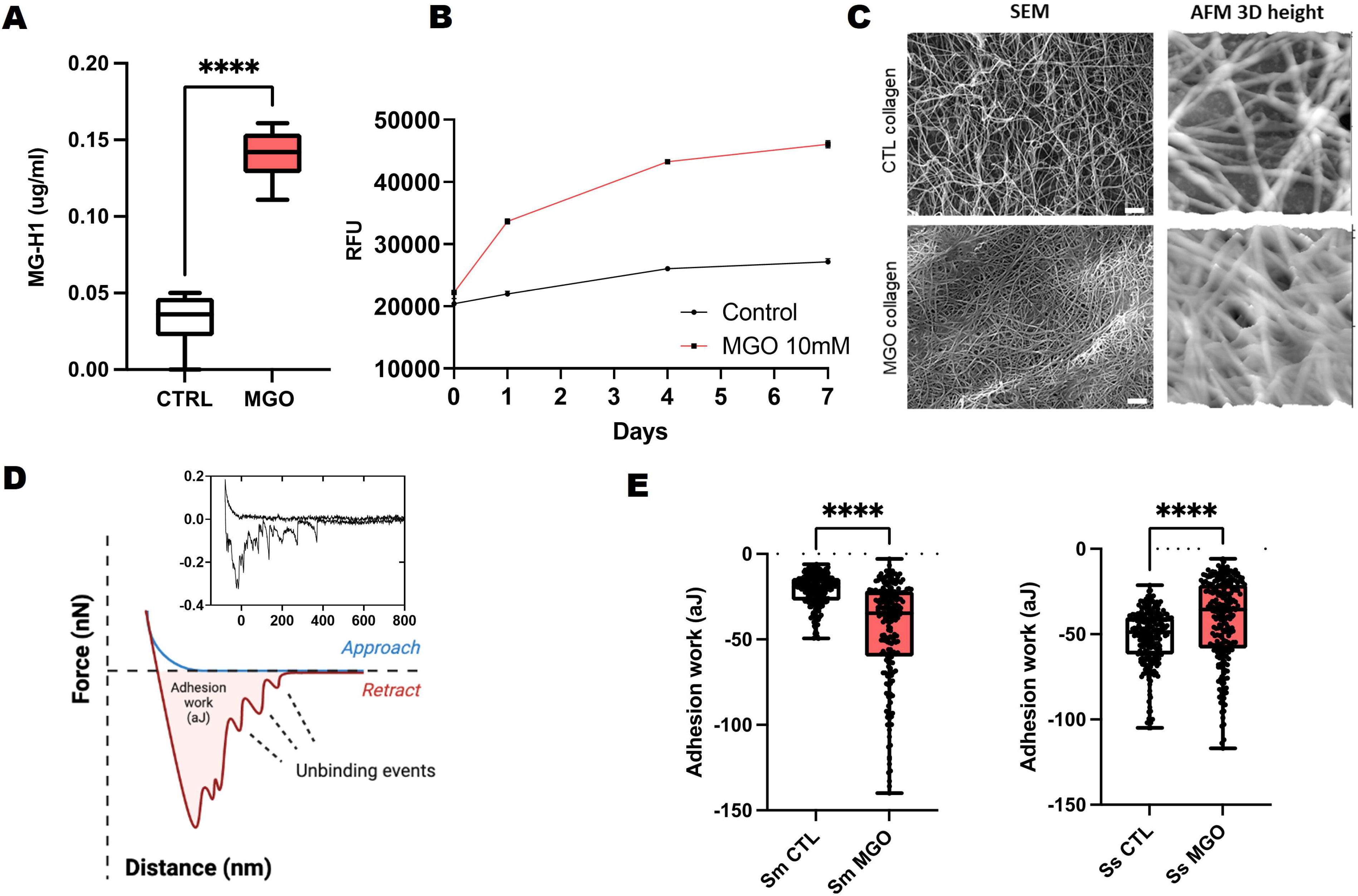
Molecular and ultrastructural characterization of in-vitro collagen substrates. (A) ELISA quantification of MG-H1 accumulation within collagen gels after 7-day incubation with MGO (****p<0.0001, paired t test). (B) Increase in collagen autofluorescence is seen for the MGO-incubated gels up to 7 days, expressed as relative fluorescence units (RFU, n=3). (C) SEM and AFM 3D reconstruction (from height scans) for both control collagen (incubated with PBS 1x) or 10mM MGO. (D) Diagrammatic representation of AFM force curves obtained between a bacterial probe and collagen substrates (Inset: experimental force curve obtained between a *Streptococcus sanguinis* probe and native collagen). (E) Work of adhesion recorded between for *Streptococcus mutans* (Sm) and *S. sanguinis* (Ss) probes on native and MGO-modified collagen substrates (****p<0.0001, Mann-Whitney test).

### Real-time nanoscale adhesion of oral streptococci onto MGO-modified collagen

With the use of non-destructive PLL-based immobilization, functionalized bacterial AFM probes were constructed to quantify the real-time adhesion of *S. mutans* and *S. sanguinis* onto native and MGO-modified collagen substrates **(Figure 1D)**. Initially, adhesion work was determined to assess the overall adhesive interaction between the probe and collagen surfaces, including the initial peak and single-unbinding events indicative of molecular bacteria-substrate interactions (Beaussart and El-Kirat-Chatel 2019). *S. mutans* was found to significantly increase its adhesion work when probed against MG-H1col, compared to native collagen **(Figure 1E)**. However, the opposite was observed for *S. sanguinis*, as MGO modification reduced its binding to collagen.

As a second step, the single-unbinding events observed between bacterial probes and collagen were explored. Therefore, the rupture force and length for individualized unbinding events (**Figure 1D**) were calculated in the Asylum Research proprietary software for all experimental conditions (**Figures 2A and 2B**). Overall, when adhering to native collagen, *S. sanguinis* demonstrated a higher number of individual molecular interactions (n=795) compared to *S. mutans* (n=265); however, MGO incorporation into collagen gels increased the number of molecular interaction sites for *S. mutans* (n=400) across the same number of total recorded probe-surface interactions. Furthermore, *S. mutans* also displayed a significant increase in its molecular binding forces when probed against the MGO-modified surface, as well as a wider spread in data towards increased interactions. For *S. sanguinis*, reduced rupture force medians for native and modified collagen were observed. No changes in rupture length **(Figure 2B)** or contour length **(Figure 2C)** were observed for either bacterial type associated with MGO incorporation into collagen.

**Figure 2:**
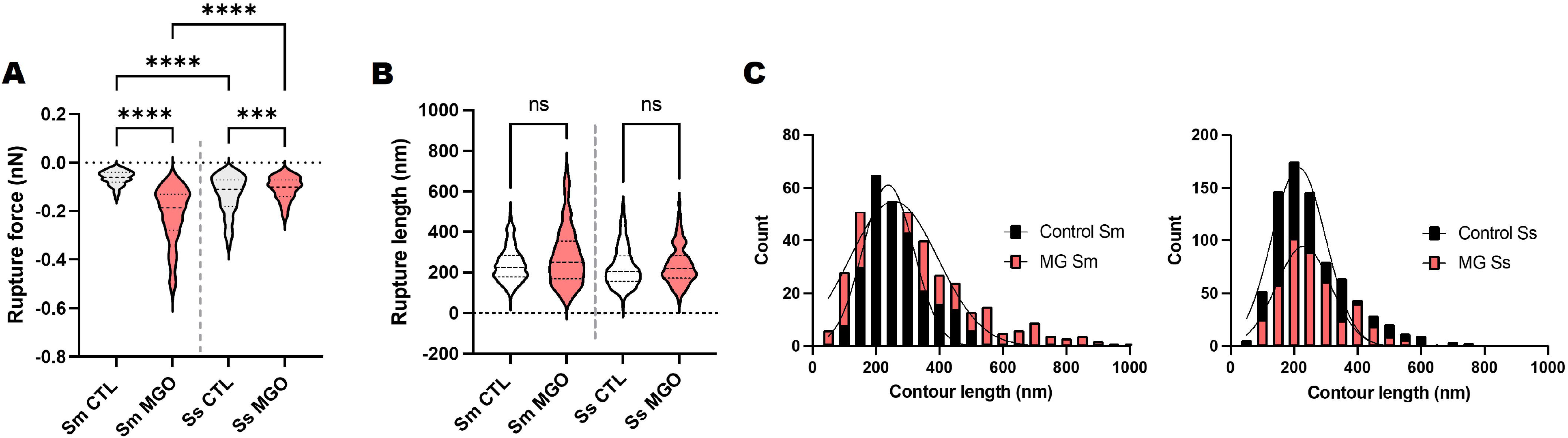
Mechanobiology of the streptococcal-collagen interaction at the sub-cellular level. (A) Violin plots for the force of individualized rupture events between streptococcal probes and collagen, demonstrating increased adhesion for *S. mutans* against MGO-modified collagen (***p<0.001; ****p<0.0001, Kruskal-Wallis test). (B) Violin plots for the rupture length of individualized unbinding events between streptococcal probes and collagen substrates (ns: non-significant). (C) Histograms (fitted to Gaussian curves) for predicted contour lengths utilizing the Worm-like chain model for both *S. mutans* and *S. sanguinis*.

To further visualize the association between rupture force and length, every recorded minor unbinding event observed between cell probes and the different collagen substrates was also plotted as a separate point on a grid **(Figure 3)**. In this format, the increase in molecular binding forces between *S. mutans* and MGO-altered becomes more obvious, compared to the interactions between the probes and native collagen substrates **(Figure 3A)**. For *S. sanguinis* no major differences were recorded between the two collagen conditions **(Figure 3B)**.

**Figure 3:**
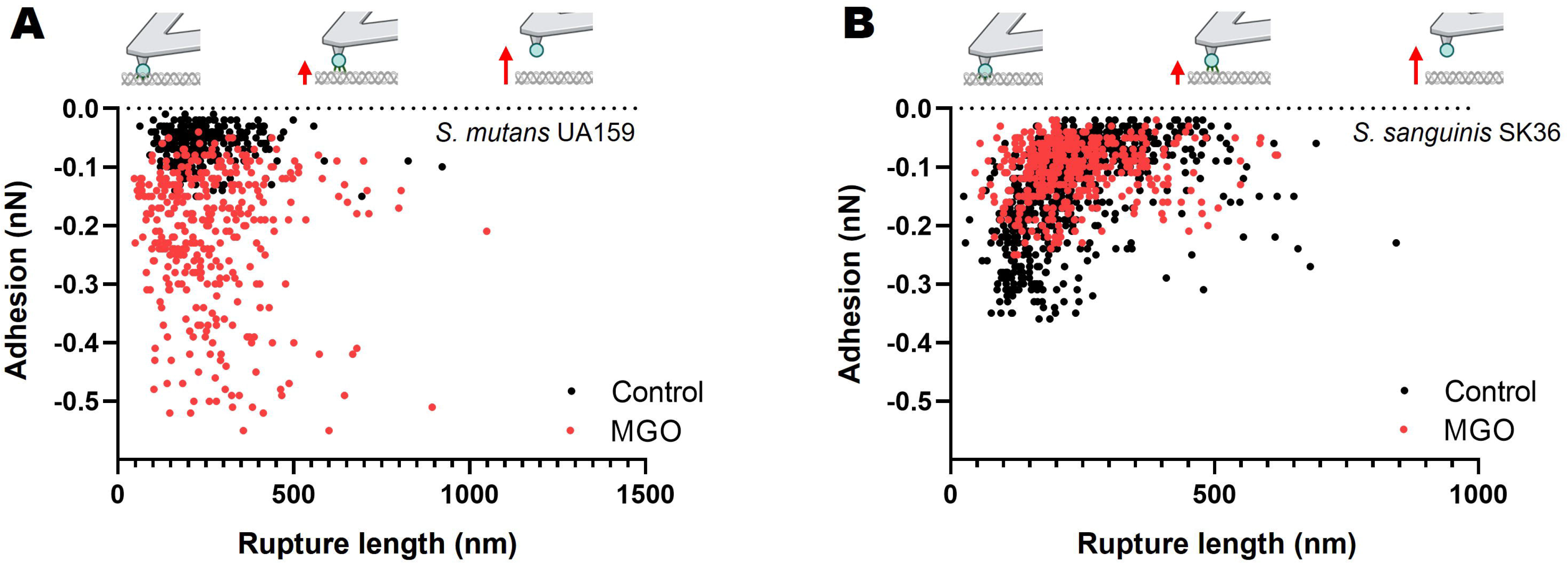
Single-unbinding event frequency and forces for *S. mutans* is modulated by MGO-modified collagen. Plot of the adhesion force vs rupture length for each single-binding event between collagen gels and (A) *S. mutans* UA 159 and (B) *S. sanguinis* SK 36. Insets: diagrammatic representation of the retraction stages between the bacterial AFM probe and collagen with respect to the X-axis. From left to right: bacteria is in complete contact with the collagen surface, initial retraction with most of the recorded single-unbinding events, probe completely removed from the collagen surface.

### Poisson analysis of streptococcal unbinding from native and MGO-modified collagen

A Poisson analysis was performed for all individual unbinding peaks according to previous approaches (Chen et al. 2011) to determine the involvement of specific and non-specific forces driving streptococcal adhesion to collagen. Our results indicated that attractive forces dominated the interaction between streptococcus and collagen for all conditions. Most importantly, there was a substantial increase in both the specific and non-specific forces between *S. mutans* and MGO-modified collagen, with the opposite being true for *S. sanguinis* **(Table 1)**.

**Table 1:** Biophysical parameters assessed for the single-unbinding events of *Streptococus sanguinis* and *Streptococcus mutans* probed against native and MGO-modified collagen gels.

|  | Number of unbindings (n) | Median single-unbinding force (nN) | Mean single-unbinding force (nN) | Poisson analysis |  |
| --- | --- | --- | --- | --- | --- |
|  |  |  |  | Short-range force (nN) | Long range-force (nN) |
| *S. mutans* UA159 control | 265 | -0.06 | -0.07 ± 0.04 | -0.04 ± 0.02 | -0.04 ± 0.03 |
| *S. mutans* UA 159 MGO | 400 | -0.19 | -0.3 ± 0.33 | -0.18 ± 0.01 | -0.13 ± 0.01 |
| *S. sanguinis* SK 36 control | 795 | -0.11 | -0.14 ± 0.1 | -0.19 ± 0.02 | -0.1 ± 0.02 |
| *S. sanguinis* SK 36 MGO | 446 | -0.1 | -0.12 ± 0.07 | -0.08 ± 0.02 | -0.07 ± 0.03 |

### Computational modeling suggests increased *S. mutans*-MGO affinity via changes in specific Cbp affinity

As a final step, we performed a protein docking simulation as a model to elucidate if MG-H1col can in fact increase *S. mutans* Cbp affinity from a theoretical standpoint. For this, we decided to utilize SpaP, a well-characterized *S. mutans* UA 159 collagen-binding adhesin that is not expressed by *S. sanguinis* SK 36 **(Supplementary Figure 1)**. Furthermore, the molecular structure for SpaP is readily available and its collagen-binding region is well documented and characterized.

Docking studies were carried out to obtain the Col/SpaP **(Figure 4A)** and MG-H1col/SpaP **(Figure 4D)** complexes. Using the HDOCK server, docking score values for Col/SpaP and MG-H1col/SpaP were −244.84 and 250.23, respectively. The best model in each case shows the interaction of SpaP with the three collagen chains (labelled as chain E, chain F and chain G). The docking score value differences can be explained based on the contact area and specific interactions established for each **complex (Figure 4B and 4E)**. Thus, in Col/SpaP, interaction from the nitrogen of hydroxyproline moiety in collagen with N1055 and Q1026 were observed. In addition, hydrogen bonds and van der Waals interactions with A1061, S1063, D1031 and Y1101 contribute to stabilizing the complex **(Figure 4B)**. Considering that an excess of MGO leads to the generation of MG-H1col via arginine modification, it is essential to mention that the S1096 and D1031 in SpaP interact with the arginine residues of Col.

**Figure 4:**
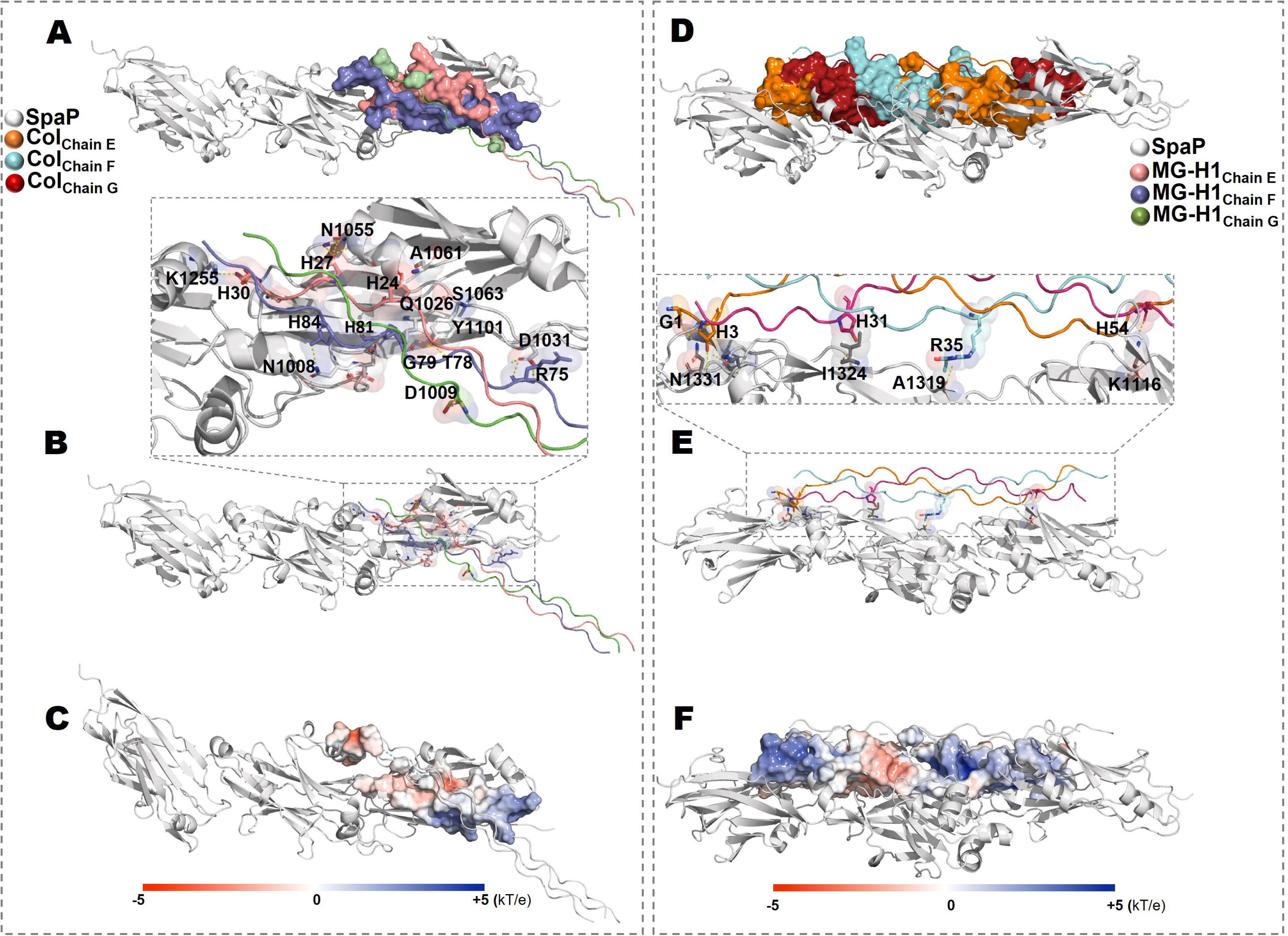
Computational modeling of the interaction between MGO-modified collagen and the biologically relevant SpaP collagen-binding protein of *S. mutans*. A) and D) the complexes Col/SpaP and Col/MG-H1 are shown. The corresponding interacting segments of Chain E-F with SpaP are displayed as surface. In B) and E) the main interactions are detailed and in C) and F) the charge distribution corresponding to the contact area is displayed (contoured from −5 (red) to +5 (blue) kT/e), using Adaptive Poisson-Boltzmann Solver (APBS) calculations.

On the other hand, the increased SpaP-MG-H1col docking score (as shown in **Figures 4D-4F**) results from a more prominent interaction region between the protein and modified collagen, leading to a more stable complex with SpaP. Furthermore, the electrostatic potential in a more extended area stabilizes the complex. The main interactions between MG-H1col/SpaP were generated with N1331, I1324, A1319, K1116, K1329, K1141, and Y1152, among others (hydrogen bonds are shown in **Figure 4E**).

## Discussion

The aging process generates significant changes in tissues including the time-dependent accumulation of AGEs. Amongst these, MGO-derived AGEs are biologically relevant in dentistry as they are known to accumulate in oral tissues such as dentin and periodontal ligament (Delle Monache et al. 2021; Schuh et al. 2022). Notably, the interaction of MGO with arginine residues in proteins including collagen induces formation of the clinically-relevant AGE MG-H1. The increase in collagen fluorescence following MGO incubation may be explained by MG-H1 carbonylation (Morimoto et al. 2017) (**Figure 1**) or via the formation of other fluorescent MGO-derived AGEs in the matrix (Ahmed et al. 2003). These results confirm previous reports that MGO-modified collagen undergoes nanomechanical and ultrastructural changes (Panwar et al. 2015; Schuh et al. 2020).

As glycation, including MGO-AGEs, is known to modulate the adhesion of host cells (Bansode et al. 2020) and resident oral bacteria (Ozer et al. 2015; Schuh et al. 2020; Śmiga et al. 2021), the effect of MGO modification on the individual biophysics of *S. mutans* and *S. sanguinis* single-cell adhesion to collagen was explored. Using AFM-based force-spectroscopy, the formation of many molecular interactions between *S. mutans* and *S. sanguinis* cells and native collagen surfaces was observed at low (0.2nN) contact forces **(Figure 2, Table 1)**. MGO-modification of collagen substrates increased the number and adhesion force of single-unbinding events for *S. mutans*, with the opposite effect being observed for *S. sanguinis*. Subsequently, no changes in rupture length or contour length were observed for either bacterial type associated with MGO incorporation into collagen **(Figure 2)**, suggesting that the increase in adhesion force between *S. mutans*-MGO is due to stronger single adhesin-collagen interactions and is not due to the recruitment of another subpopulation of adhesins (e.g., with a different molecular length). In literature, similar minor rupture lengths of mainly <500nm were observed for AFM experiments with *S. mutans* and *S. sanguinis* against a range of surfaces (Hwang et al. 2015; Aguayo et al. 2016; Wang et al. 2020). Furthermore, force-spectroscopy experiments with isolated the *S. mutans* collagen-binding SpaP demonstrate similar rupture lengths and forces (Heim et al. 2015; Sullan et al. 2015). Therefore, the present results are within the expected values for streptococci probed against soft, compliant biological surfaces at reduced probe loading forces (Wan et al. 2020).

There can be many potential ways to explain the increase in *S. mutans* binding to MGO-modified collagen seen in our system, including geometrical factors and changes in the molecular conformation of the collagen molecule following AGE incorporation. In this work, in-silico approaches employing the relevant *S. mutans* UA159 SpaP Cbp (Ajdić et al. 2002) suggest that MGO modification of type-I collagen increases the docking affinity between this adhesin and the substrate. The present simulations show that the generation of MG-H1 within the region of collagen known to interact with SpaP increases the contact area and docking score for the collagen-SpaP complex **(Figure 4)**. This data supports the bacterial probe force-spectroscopy results showing increased binding forces between MGO-modified collagen and *S. mutans* UA 159 but not *S. sanguinis* SK 36, particularly as SpaP is not expressed by *S. sanguinis* **(Supplementary Figure 1)**. The molecular length of SpaP is also consistent with the rupture and contour lengths probed with AFM in-vitro in *S. mutans* **(Figure 2)**; nevertheless, oral streptococci utilize a range of adhesins and other structures to attach to surfaces, and further work should elucidate the role of other relevant Cbps in the binding to AGE-modified collagen substrates (Álvarez et al. 2021).

Furthermore, recent work has shown that another important oral pathogen, *P. gingivalis*, attaches more robustly to glycated collagen surfaces (Śmiga et al. 2021), and that AGE accumulation in tissues promotes pathologic bacterial adhesion in the urinary tract (Ozer et al. 2015). Altogether, these reports suggest that adhesion to glycated tissues may be an important virulence factor across different strains of disease-generating bacteria from our microbiome, and that it may not be particular to *S. mutans* UA 159 or oral streptococci. This is biologically relevant as collagen is one of the most relevant substrates for initial attachment and as such, plays an essential role in pathogenic adhesion by early colonizers in teeth and remote tissues (Singh et al. 2012; Avilés-Reyes et al. 2017). The implications of aging-associated collagen changes on oral bacterial invasion and translocation to other organs remains to be explored. Also, additional studies should explore the possible effect of other relevant collagen AGEs – such as pentosidine – on bacterial adhesion and early-biofilm progression, as well as their potential impact on surface colonization by bacterial aggregates in the oral cavity in the context of health and disease (Simon-Soro et al. 2022).

## Conclusion

Higher *S. mutans* adhesion onto MGO-modified collagen is mediated by increasing the number and force of specific cell-substrate interactions at the sub-cellular level, as explored with living bacterial AFM probes and bacterial force-spectroscopy. Both short-range and long-range adhesive forces were increased for the *S. mutans*-MGO collagen interaction. Neither of these effects was observed for *S. sanguinis* utilizing the same approach, suggesting that this effect is strain specific and associated to the particular bacterial adhesins expressed by *S. mutans* UA 159. Computational in-silico approaches using the relevant *S. mutans* UA159 adhesin SpaP suggest an increase in the docking affinity between this adhesin and collagen molecules following MGO incorporation.

## Acknowledgments

This work was funded by the ANID FONDECYT Grants #11180101 and #1220804, and the Millennium Science Initiative #P10-035F. Author contributions: CLS contributed to data acquisition, analysis, and interpretation, and critically revised the manuscript. PTL, LHG, and LR contributed to data acquisition and critically revised the manuscript. AR and AF contributed to data acquisition, analysis, and interpretation, and critically revised the manuscript. NPB and LB contributed to design, data interpretation, and critically revised the manuscript. CS contributed to design, data acquisition and interpretation, and critically revised the manuscript. SA contributed to conception, design, data acquisition and interpretation, drafted and critically revised the manuscript, and acquired funding for the study. All authors gave their final approval and agree to be accountable for all aspects of the work.

## Notes

### Competing Interest Statement

The authors have declared no competing interest.

## References

Aguayo S, Donos N, Spratt D, Bozec L. 2016. Probing the nanoadhesion of Streptococcus sanguinis to titanium implant surfaces by atomic force microscopy. Int J Nanomedicine. 11:1443–50.

Ahmed N, Thornalley PJ, Dawczynski J, Franke S, Strobel J, Stein G, Haik GM. 2003. Methylglyoxal-Derived Hydroimidazolone Advanced Glycation End-Products of Human Lens Proteins. Invest Ophthalmol Vis Sci. 44(12):5287–5292.

Ahmed T, Nash A, Clark KE, Ghibaudo M, de Leeuw NH, Potter A, Stratton R, Birch HL, Enea Casse R, Bozec L. 2017. Combining nano-physical and computational investigations to understand the nature of “aging” in dermal collagen. Int J Nanomedicine. 12:3303–3314.

Ajdić D, McShan WM, McLaughlin RE, Savić G, Chang J, Carson MB, Primeaux C, Tian R, Kenton S, Jia H, et al. 2002. Genome sequence of Streptococcus mutans UA159, a cariogenic dental pathogen. Proceedings of the National Academy of Sciences. 99(22):14434–14439.

Álvarez S, Leiva-Sabadini C, Schuh CMAP, Aguayo S. 2021. Bacterial adhesion to collagens: implications for biofilm formation and disease progression in the oral cavity. Crit Rev Microbiol.:1–13.

Avilés-Reyes A, Miller JH, Lemos JA, Abranches J. 2017. Collagen-binding proteins of Streptococcus mutans and related streptococci. Mol Oral Microbiol. 32(2):89–106.

Bailey AJ, Paul RG, Knott L. 1998. Mechanisms of maturation and ageing of collagen. Mech Ageing Dev. 106(1):1–56.

Baker NA, Sept D, Joseph S, Holst MJ, McCammon JA. 2001. Electrostatics of nanosystems: Application to microtubules and the ribosome. Proceedings of the National Academy of Sciences. 98(18):10037–10041.

Bansode S, Bashtanova U, Li R, Clark J, Müller KH, Puszkarska A, Goldberga I, Chetwood HH, Reid DG, Colwell LJ, et al. 2020. Glycation changes molecular organization and charge distribution in type I collagen fibrils. Sci Rep. 10(1):3397.

Beaussart A, El-Kirat-Chatel S. 2019. Microbial adhesion and ultrastructure from the single-molecule to the single-cell levels by Atomic Force Microscopy. The Cell Surface. 5:100031.

Berne C, Ellison CK, Ducret A, Brun Y V. 2018. Bacterial adhesion at the single-cell level. Nat Rev Microbiol. 16(10):616–627.

Chen Y, Busscher HJ, van der Mei HC, Norde W. 2011. Statistical Analysis of Long- and Short-Range Forces Involved in Bacterial Adhesion to Substratum Surfaces as Measured Using Atomic Force Microscopy. Appl Environ Microbiol. 77(15):5065–5070.

Delle Monache S, Pulcini F, Frosini R, Mattei V, Talesa VN, Antognelli C. 2021. Methylglyoxal-Dependent Glycative Stress Is Prevented by the Natural Antioxidant Oleuropein in Human Dental Pulp Stem Cells through Nrf2/Glo1 Pathway. Antioxidants. 10(5).

Egaña-Gorroño L, López-Díez R, Yepuri G, Ramirez LS, Reverdatto S, Gugger PF, Shekhtman A, Ramasamy R, Schmidt AM. 2020. Receptor for Advanced Glycation End Products (RAGE) and Mechanisms and Therapeutic Opportunities in Diabetes and Cardiovascular Disease: Insights From Human Subjects and Animal Models. Frontiers in Cardiovascular Medicine. 7:37.

Gkogkolou P, Böhm M. 2012. Advanced glycation end products: Key players in skin aging? Dermatoendocrinol. 4(3):259–270.

Gurav AN. 2013. Advanced Glycation End Products: A Link Between Periodontitis and Diabetes Mellitus? Curr Diabetes Rev. 9(5):355–361.

Heim KP, Sullan RMA, Crowley PJ, El-Kirat-Chatel S, Beaussart A, Tang W, Besingi R, Dufrene YF, Brady LJ. 2015. Identification of a Supramolecular Functional Architecture of Streptococcus mutans Adhesin P1 on the Bacterial Cell Surface*. Journal of Biological Chemistry. 290(14):9002–9019.

Hojo K, Nagaoka S, Ohshima T, Maeda N. 2009. Bacterial interactions in dental biofilm development. J Dent Res. 88(11):982–90.

Hwang G, Marsh G, Gao L, Waugh R, Koo H. 2015. Binding Force Dynamics of Streptococcus mutans–glucosyltransferase B to Candida albicans. J Dent Res. 94(9):1310–1317.

Kontis V, Bennett JE, Mathers CD, Li G, Foreman K, Ezzati M. 2017. Future life expectancy in 35 industrialised countries: projections with a Bayesian model ensemble. Lancet. 389(10076):1323–1335.

Morimoto H, Gu L, Zeng H, Maeda K. 2017. Amino Carbonylation of Epidermal Basement Membrane Inhibits Epidermal Cell Function and Is Suppressed by Methylparaben. Cosmetics. 4(4).

Murray Thomson W. 2014. Epidemiology of oral health conditions in older people. Gerodontology. 31(s1):9–16.

Nass N, Bartling B, Santos AN, Scheubel RJ, Börgermann J, Silber RE, Simm A. 2007. Advanced glycation end products, diabetes and ageing. Z Gerontol Geriatr. 40(5):349–356.

Nomura R, Nakano K, Taniguchi N, Lapirattanakul J, Nemoto H, Grönroos L, Alaluusua S, Ooshima T. 2009. Molecular and clinical analyses of the gene encoding the collagen-binding adhesin of Streptococcus mutans. J Med Microbiol. 58(4):469–475.

Ozer A, Altuntas CZ, Izgi K, Bicer F, Hultgren SJ, Liu G, Daneshgari F. 2015. Advanced glycation end products facilitate bacterial adherence in urinary tract infection in diabetic mice. Pathog Dis. 73(5):ftu004.

Panwar P, Lamour G, Mackenzie NCW, Yang H, Ko F, Li H, Brömme D. 2015. Changes in Structural-Mechanical Properties and Degradability of Collagen during Aging-associated Modifications *. Journal of Biological Chemistry. 290(38):23291–23306.

Pettersen EF, Goddard TD, Huang CC, Couch GS, Greenblatt DM, Meng EC, Ferrin TE. 2004. UCSF Chimera—A visualization system for exploratory research and analysis. J Comput Chem. 25(13):1605–1612.

Schuh CMAP, Benso B, Naulin PA, Barrera NP, Bozec L, Aguayo S. 2020. Modulatory Effect of Glycated Collagen on Oral Streptococcal Nanoadhesion. J Dent Res. 100(1):82–89.

Schuh CMAP, Leiva-Sabadini C, Huang S, Barrera NP, Bozec L, Aguayo S. 2022. Nanomechanical and Molecular Characterization of Aging in Dentinal Collagen. J Dent Res. 101(7):840–847.

Semba RD, Nicklett EJ, Ferrucci L. 2010. Does accumulation of advanced glycation end products contribute to the aging phenotype? J Gerontol A Biol Sci Med Sci. 65(9):963–975.

Simon-Soro A, Ren Z, Krom BP, Hoogenkamp MA, Cabello-Yeves PJ, Daniel SG, Bittinger K, Tomas I, Koo H, Mira A. 2022. Polymicrobial Aggregates in Human Saliva Build the Oral Biofilm. mBio. 13(1):e0013122–e0013122.

Singh B, Fleury C, Jalalvand F, Riesbeck K. 2012. Human pathogens utilize host extracellular matrix proteins laminin and collagen for adhesion and invasion of the host. FEMS Microbiol Rev. 36(6):1122–1180.

Śmiga M, Smalley JW, Ślęzak P, Brown JL, Siemińska K, Jenkins RE, Yates EA, Olczak T. 2021. Glycation of Host Proteins Increases Pathogenic Potential of Porphyromonas gingivalis. Int J Mol Sci. 22(21).

Sullan RMA, Li JK, Crowley PJ, Brady LJ, Dufrêne YF. 2015. Binding Forces of Streptococcus mutans P1 Adhesin. ACS Nano. 9(2):1448–1460.

Takahashi N, Nyvad B. 2016. Ecological Hypothesis of Dentin and Root Caries. Caries Res. 50(4):422–431.

Wan SX, Tian J, Liu Y, Dhall A, Koo H, Hwang G. 2020. Cross-Kingdom Cell-to-Cell Interactions in Cariogenic Biofilm Initiation. J Dent Res. 100(1):74–81.

Wang R, Deng L, Lei Z, Wu P, Wang Y, Hao L, Li T, Jiang L. 2020. Nanoscale adhesion forces of glucosyltransferase B and C genes regulated Streptococcal mutans probed by AFM. Mol Oral Microbiol. 35(2):49–55.

Wetzels S, Wouters K, Schalkwijk CG, Vanmierlo T, Hendriks JJA. 2017. Methylglyoxal-Derived Advanced Glycation Endproducts in Multiple Sclerosis. Int J Mol Sci. 18(2):421.

Yan Y, Tao H, He J, Huang S-Y. 2020. The HDOCK server for integrated protein– protein docking. Nat Protoc. 15(5):1829–1852.

Yukari I, Toru T, Shinya K, Rie M, Mikari A, Yukie S, Yuki S, Kunio I, Ichiro T, Yoshihisa Y, et al. 2021. Identification of Initial Colonizing Bacteria in Dental Plaques from Young Adults Using Full-Length 16S rRNA Gene Sequencing. mSystems. 4(5):e00360-19.

