## Supplementary data 1 for "Nanoscale dynamics of streptococcal adhesion to AGE-modified collagen"

**SpaP detection with conventional PCR**

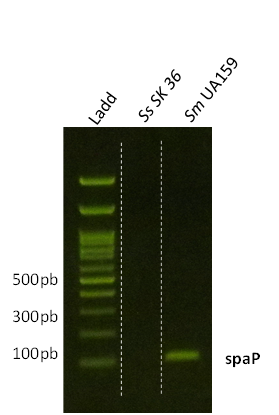
From overnight cultures of *Streptococcus mutans* UA 159 and *Streptococcus sanguinis* SK 36, individual colonies were resuspended in distilled water and heated to 95°C for 10 minutes to lyse bacterial cells and release DNA. Samples were then centrifuged, and the DNA-containing supernatants were processed with a commercially-available kit (Go Taq®, Promega, USA) utilizing custom-made primers for SpaP according to previously published protocols (Park et al. 2019). A 1000bp DNA ladder was used to determine fragment sizes, and samples were loaded into an electrophoresis gel with 6x loading dye (ThermoFisher, USA), and visualized with the SYBR Safe DNA gel stain (Invitrogen, USA).

**Supplementary figure 1**: (A) *Streptococcus mutans* UA159 expresses the relevant collagen-binding protein SpaP, which is absent in *Streptococcus sanguinis* SK 36.

**Reference**:

Park, B.-I., Kim, B.-S., Kim, K.-J., & You, Y.-O. (2019). Sabinene suppresses growth, biofilm formation, and adhesion of Streptococcus mutans by inhibiting cariogenic virulence factors. *Journal of Oral Microbiology*, *11*(1), 1632101. https://doi.org/10.1080/20002297.2019.1632101
